# *In vitro* fertilisation procedure assisted with computer vision models for organic Senegalese sole (*Solea senegalensis*) culture

**DOI:** 10.64898/2026.02.26.707220

**Authors:** Abdul Qadir, Sandra Salcedo Martinez, Francesc Serratosa, Neil Duncan

**Affiliations:** IRTA, Aquaculture, Centre de La Ràpita, 43540 La Ràpita, Catalonia, Spain; Universitat Rovira i Virgili, Department of Computer Engineering and Mathematics (DEIM), Tarragona, Spain

**Author notes:** Corresponding authors: Neil Duncan, and Francesc Serratosa.

**Keywords:** Aquaculture, Reproductive behavior, Senegalese Sole, Tracking, Artificial intelligence, Deep learning, Convolutional neural networks

## Abstract

Reproductive dysfunction remains a major challenge for Senegalese sole (*Solea senegalensis*) aquaculture. Hormone-induced ovulation and *in vitro* fertilisation (IVF) are currently used to overcome the absence of natural courtship behaviours in captivity. This study investigates the feasibility of hormone-free IVF and behaviour-based prediction of ovulation as alternative strategies to enhance reproductive outcomes. We selected males using computer-assisted sperm analysis to assess sperm motility and quality for IVF trials. IVF trials were conducted using selected males and naturally ovulated eggs collected from females during evening hours across six experimental nights in two groups. Fish behaviour was continuously recorded using underwater cameras, and a convolutional neural network was developed to automatically detect Rest the Head (RTH) and Locomotor Activity (LA) behaviours. These behavioural counts, together with timing information, were used as features to train a logistic regression model for predicting ovulation events. Hormone-free IVF achieved fertilization rates up to 44% with 18% hatching success, producing viable larvae without hormonal intervention. Both groups showed significantly elevated RTH and LA during ovulation nights compared to non-ovulation nights, with peak activity occurring between 18:00–19:00 hours. The behavioural prediction model correctly identified ovulation with 82–85% accuracy and an area under the curve of 0.95. These findings demonstrate that sperm-quality-based male selection combined with automated behaviour analysis provides a practical, non-invasive approach for hormone-free reproduction in organic flatfish aquaculture.

## 1. Introduction

Senegalese sole (*Solea senegalensis*) has become an important species in European aquaculture, particularly in Spain, due to its disease resistance, favourable growth rates, and high market demand. These attributes, coupled with advancements in closed-cycle reproduction and recirculating aquaculture systems (RAS), have facilitated its culture and contributed to the industry’s sustainability (Anguis and Cañavate, 2005; APROMAR, 2022, 2019; Morais et al., 2016).

However, large-scale production of Senegalese sole faces numerous challenges in reproduction. A major issue in captive broodstocks is reproductive failure, which has been linked to the absence of natural courtship behaviours and the non-fertilisation of eggs (Duncan et al., 2019; Martín et al., 2020). Spawning attempts in captive populations frequently fail to produce viable offspring, highlighting the need for an improved understanding and management of reproductive behaviours in aquaculture settings (Carazo et al., 2016; Martín et al., 2020). As natural spawning is not observed in cultured broodstock, *in vitro* fertilisation (IVF) has been implemented and is the only reliable method to produce fertilised eggs from cultured breeders. However, successful IVF depends on accurate timing of female ovulation, which is traditionally achieved through hormone treatments with gonadotropin-releasing hormone agonists (GnRHa) (Rasines et al., 2012; Ramos-Júdez et al., 2021). These hormones are used to induce and synchronize ovulation and to control its timing, allowing for the timely collection of eggs for IVF. Nevertheless, the use of hormones is prohibited in organic aquaculture, which follows ecological and ethical standards (European Commission, 2021a). As a result, there is a need to develop hormone-free methods to obtain viable ovulated eggs through methods such as predicting natural ovulation. Such approaches would enable sustainable reproduction in compliance with organic aquaculture principles, helping the industry reduce its dependence on hormonal treatments and improving the ethical and environmental profile of aquaculture systems. Behavioural studies have shown that certain courtship behaviours in Senegalese sole are closely linked to reproductive readiness and spawning events (Duncan et al., 2019; Fatsini et al., 2020; Qadir et al., 2025). Monitoring these behaviours presents a promising non-invasive alternative for anticipating ovulation timing in the absence of authorisation to use hormonal factors.

Recent advances in machine learning have considerable potential in aquaculture by optimizing breeding strategies and improving reproductive management (Kotsiantis et al., 2006; Shinde and Shah, 2018). Machine learning algorithms can detect patterns in complex biological data, including fish behaviour, that may not be apparent through traditional observation methods. For example, deep learning models combining ResNet50 with long short-term memory networks have successfully recognized five reproductive behaviours during fish spawning, including chasing and spawning behaviours, achieving 98.52% accuracy (Du et al., 2022). These applications demonstrate the potential of machine learning to automate behavioural monitoring in aquaculture settings.

The aim of this article is twofold: first, to demonstrate that naturally ovulated eggs can be successfully used for IVF without hormonal intervention, providing a proof of concept for hormone-free reproduction in organic aquaculture; and second, to develop a machine learning model that predicts ovulation timing from automated behavioural monitoring. The model would enable timely gamete collection for commercial-scale IVF utilization. The rest of the article is organised as follows. In the next section, the experimental setting and the process of obtaining the data are explained. Moreover, the automated system to detect the reproductive behaviours are detailed. Then, Section 3 explains the obtained results that validate our system and demonstrate ovulated eggs can be successfully used for IVF without hormonal intervention. Section 4 and Section 5 discusses the experiments and concludes the article, respectively.

## 2. Material and methodology

### 2.1 Experimental settings

The experimental data were collected in IRTA La Ràpita from March 14 to May 22, 2024. Two experimental groups were housed in 10 m³ fiberglass tanks, each equipped with an IRTAmar® recirculation system at IRTA La Ràpita. The first group Mix1 consisted of 17 fish, including 5 cultured males, 2 cultured females, 8 wild males, and 2 wild females (mean weight = 760 ± 416 g, range: 342-1708 g). The second group, Mix2, included 16 fish, comprising 1 cultured male, 5 cultured females, 7 wild males, and 3 wild females (mean weight = 673 ± 541 g, range: 151-2336 g). On average, the cultured and wild fish had been in captivity for approximately 6 to 7 years.

Both tanks were maintained under natural environmental conditions for one year prior to the experiment. During the experimental period, water temperature ranged naturally from 9 to 20°C, and photoperiod varied naturally from 9 to 14 hours of light per day. To stimulate spawning, temperature was manipulated weekly once it reached 18°C: Monday through Thursday at 16 ± 1°C, and Friday through Sunday at 18 ± 1°C (Martín et al., 2014; Fatsini, 2017). Fish were fed four days per week with 0.75% wet feed (polychaetes and mussels) and 0.55% dry feed (balanced commercial diet) relative to total biomass. Each fish was individually tagged with a passive integrated transponder (PIT tag: ID-100 Unique, Trovan-Zeus, Madrid, Spain) for identification.

### 2.2 Spawning and egg collection

During the spawning season (March to May 2024), spawning was monitored and eggs were collected daily from both experimental groups. Reproductive behaviour and fertilisation success were closely observed to assess overall spawning activity. Water temperature was recorded throughout the study period. Individual egg collectors were installed at the surface water outflow of each tank to capture buoyant eggs released by females during spawning events. Each morning at 09:00 h, collectors were inspected to ensure timely retrieval of spawned eggs. For each spawn, the following parameters were assessed: total egg volume (mL), volume of floating (viable) and non-floating (non-viable) eggs (mL), total number of floating eggs, fertilisation rate (%), hatching rate (%), and total number of larvae. Fertilisation success and developmental stage were determined by examining a subsample of approximately 120 eggs under a stereomicroscope at the time of collection and again 24 hours later. The floating egg fraction was measured using a graduated cylinder and transferred to an incubator (31 cm diameter × 33 cm depth, approximately 25 L total volume) maintained at ambient temperature and photoperiod with continuous water flow.

Larvae hatched after an incubation period of 36-48 hours. Hatching rate was assessed by counting eggs and larvae from three 100 mL subsamples taken from the thoroughly mixed incubator contents. The collection of eggs and evaluation of egg quality parameters confirmed ovulation occurrence and enabled evaluation of overall reproductive performance.

### 2.3 Assessing ovulation and IVF

For IVF procedures, gamete collection was conducted between 14:00 and 24:00 h during March and April 2024, coinciding with the natural spawning season to ensure availability of ovulated eggs. Sperm collection from males was performed from 14:00 to 18:00 h. Sperm was obtained from individual males by first locating the testes through gentle abdominal palpation. Once identified, gentle abdominal pressure was applied toward the urogenital pore to massage the testes and express sperm. Sperm was collected directly into a clean 1 mL syringe containing 0.3 mL of modified Leibovitz. As demonstrated by González-López et al. (2020), avoiding urine contamination during stripping is nearly impossible, but high numbers of motile spermatozoa can still be obtained from samples containing urine. The massage and stripping process was repeated to obtain the complete sample. The volume of collected sperm was measured to an accuracy of 10 μL using the syringe graduations, then transferred to a 2 mL microcentrifuge tube and immediately diluted 1:3 (v/v) with modified Leibovitz medium. Each sample was labelled with the fish’s PIT tag ID and collection date, then placed in a rack over crushed ice and stored at 4°C until CASA analysis or fertilisation.

For quantitative motility assessment, sperm activation was performed immediately before analysis. Depending on cell concentration, 10 or 20 μL of diluted sperm was added to 250 μL of activation solution (natural seawater supplemented with 3% w/v Bovine Serum Albumin (BSA; Sigma-Aldrich, Spain)) and gently mixed. Immediately, 2 μL of activated sperm was transferred to a sperm counting chamber pre-positioned under the microscope.

Sperm motility and kinetics were assessed within one hour of collection using a Computer-Assisted Sperm Analysis (CASA) system. The spermatozoa were recorded within 15 seconds of activation for a duration of 2 seconds using IC Capture software and a GigE digital camera (The Imaging Source GmbH) mounted on a Nikon Eclipse Ci microscope integrated with the CASA system (Microoptics, Barcelona, Spain). For each fish, three independent sperm samples were analysed. The CASA software automatically assessed the sperm quality parameters: cell concentration (cells/mL), motility (%), curvilinear velocity (VCL, μm/s), average path velocity (VAP, μm/s), straight-line velocity (VSL, μm/s), straightness (STR %), linearity (LIN %), wobble (WOB %), amplitude of lateral head displacement (ALH, μm) and beat cross frequency (BCF, Hz).

In the evening, between 22:00 and 24:00 h, females from both experimental groups, Mix1 and Mix2, were checked for signs of ovulation on six different nights during the spawning season. Prior to the egg stripping process, each female was anaesthetised in a 30-L seawater bath containing 60 µg L⁻¹ of tricaine methanesulfonate (MS-222; Acros-Organic, New Jersey, USA). Once anaesthetised, the fish tag ID, body weight (in grams), and total length (in centimetres) were registered, and each fish was checked for the presence of eggs. Females with the presence of eggs were washed with clean water, blotted dry and the ovulated eggs were extracted by gently applying abdominal pressure to collect the eggs in a 500 mL beaker. The total volume of collected eggs was measured in mL. The eggs were then distributed into 100 mL beakers, with 0.5 mL of eggs added to each beaker. Immediately after the eggs were placed in the beaker, a predetermined volume of sperm was added to ensure a minimum of 4 million motile spermatozoa per beaker (approximately 13,000 to 28,000 motile spermatozoa per egg, depending on egg density). Immediately following sperm addition, 10 mL of clean seawater was added, and the beaker was gently agitated to mix the eggs, sperm and water, ensuring sperm activation to fertilise the eggs. After 2-5 minutes, the beaker was filled with clean seawater. Each beaker was labelled to indicate the combination of female and male sperm used in fertilisation. For each male-female pairing, three replicate fertilisations were made. The remaining eggs were also fertilised in a large-scale fertilisation by mixing with sperm samples from different males and clean seawater. The beakers containing the eggs, sperm, and seawater were then stored at a temperature of 16°C in a thermostat-controlled laboratory incubator, while large scale fertilisations were placed in a flow-through fish egg incubator (31 cm diameter × 33 cm depth, approximately 25 L total volume).

### 2.4 Reproductive Behaviour

Reproductive behaviour in Senegalese sole plays an important role in predicting ovulation, which precedes and enables successful spawning. Analysis of reproductive behaviour was focused on two behaviours that we found were indicators of spawning (Qadir et al., 2025):

- **Rest the Head (RTH)**: This behaviour occurs when one fish rests its head on another fish’s body, typically observed as part of courtship interactions. An increase in this behaviour has been associated with mating courtships and is a critical indicator of the fish’s readiness for spawning.
- **Locomotor Activity (LA)**: This behaviour is defined as occurring when the distance travelled by a tracked fish between frames exceeds a speed threshold of 4 pixels/frame over a minimum of 48 frames (equivalent to approximately 2 seconds at 24 FPS). It represents dynamic behaviour that has been shown to increase with reproductive engagement, as fish engage in more active and coordinated behavioural patterns during courtship and spawning activities.

### 2.5 Video and Image Collection

Two tanks, Mix1 and Mix2, were equipped with cameras (Model SW-SK5-2D5X10-VRC, Barlus, Shenzhen, China) capable of recording in low-light conditions using red lights to facilitate nighttime recording. The cameras were positioned just below the water’s surface at the corner of each tank to record daily fish activity during the spawning season. The cameras were connected to a computer for recording videos and video streaming. Fish activity was monitored and recorded continuously (1920 × 1080 pixels per frame and 24 frames per second) for six hours daily (14:00-20:00 h) from March 18 to May 6 in both tanks.

From these recorded videos, six spawning and six non-spawning nights were selected to generate images dataset for CNN model training. Images were categorised into two behaviours: the previously described RTH behaviour, and instances when a fish was alone, which was named Fish Alone (FA) and identified the presence of fish that were not engaged in the RTH behaviour. From each night, a two-hour period from 18:00 to 20:00 was chosen specifically to capture a mix of behaviours and maximize the number of instances of both RTH and FA for the dataset, resulting in a total of 24 hours of video data (12 nights × 2 hours/night). Instances of RTH and FA with edges detected by the AI tool RoboFlow are shown in Figure 1.

**Figure 1.**
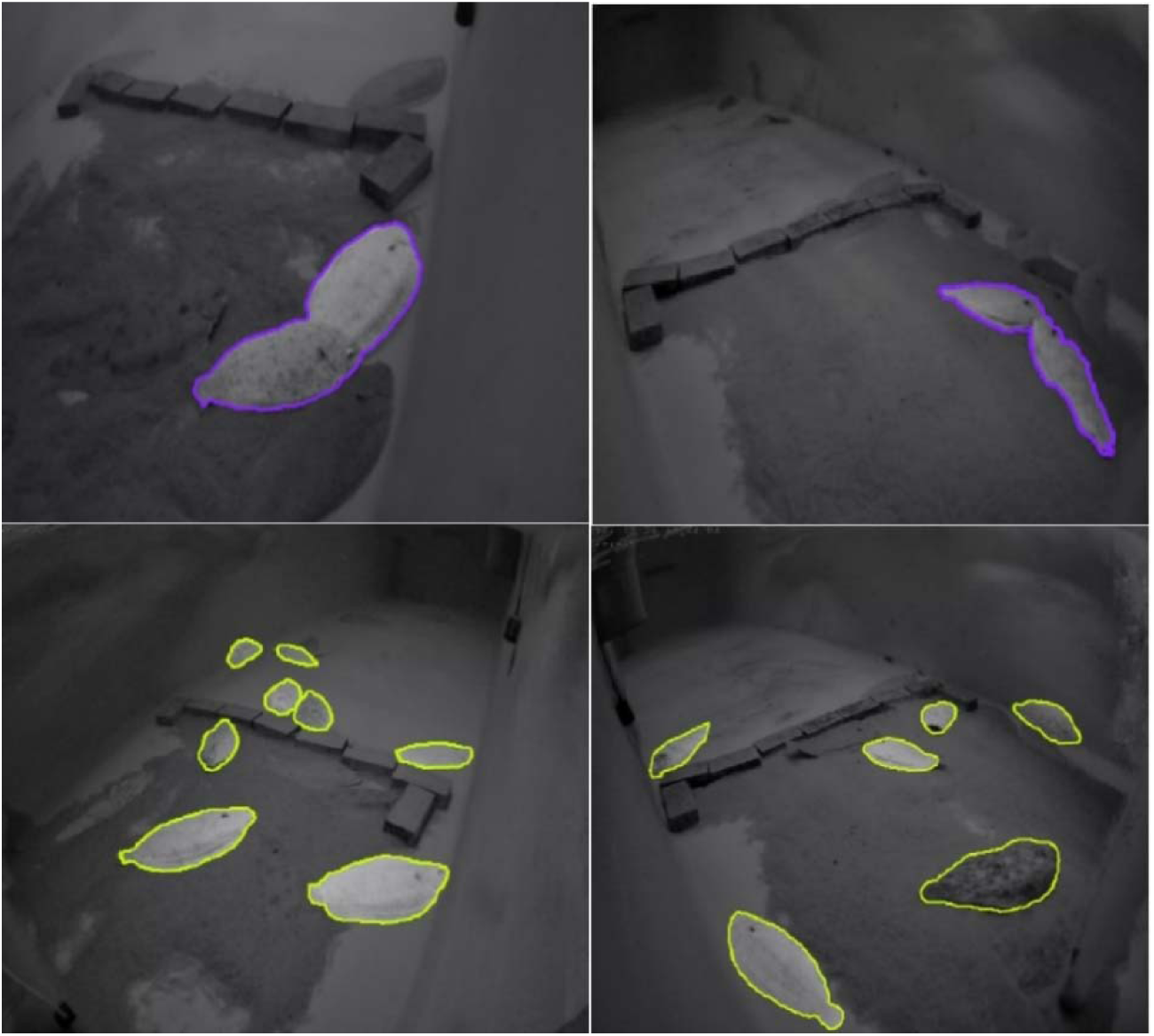
Upper images: appearance of RTH behaviour. Lower images: Appearance of several FA. Edges were detected by the AI tool RoboFlow.

A total of 2,000 images were manually selected from these 24 hours of video. Of these, 1,000 images containing RTH behaviour and another 1,000 for fish being alone (FA). Moreover, data augmentation transformations were applied to enhance this initial dataset. Among several augmentation techniques, these included flipping, 90-degree rotation, random rotation, random crop, random shear, blur, and exposure adjustments. A final database of 5,000 images was generated, which were used for training the computer vision system. Images containing RTH behaviour were used to train this behaviour and images containing FA were used to train LA.

### 2.6 Behavioural Analysis

For behavioural analysis, recorded videos were taken into consideration from a total of 18 nights for group Mix1 and 21 nights for group Mix2. Nights were classified into five categories based on reproductive outcomes.

Ovulation nights with fertilised eggs were nights followed by collection of naturally fertilised eggs, confirming successful ovulation and natural mating (Mix1: n = 2; Mix2: n = 3). Ovulation nights with unfertilised eggs were nights followed by collection of unfertilised eggs, indicating ovulation occurred but natural fertilisation failed (Mix1: n = 4; Mix2: n = 6). No ovulation nights were nights with no eggs collected the following morning (Mix1: n = 6; Mix2: n = 6). Ovulation nights with IVF fertilised eggs were nights that females were checked for ovulation, eggs were collected for IVF, and fertilisation was successful (Mix1: n = 0; Mix2: n = 4). Ovulation nights with IVF unfertilised eggs were nights that females were checked for ovulation, eggs were collected for IVF, but fertilisation was unsuccessful (Mix1: n = 6; Mix2: n = 2).

The trained CNN model was used to quantify RTH and LA behaviours at five hourly time points (16:00, 17:00, 18:00, 19:00, and 20:00 h) from each selected night. These behavioural counts were then used as input features for the ovulation prediction model described in Section 2.7.

### 2.7 Ovulation predictor model

The model to detect behaviours and predict ovulation followed the flow schema in Figure 2. The model starts with a real-time input video during the afternoon and computes a probability of ovulation for the night. Considering this probability that evolves during the afternoon, the researcher or aquaculture technician takes actions such as modifying the environment conditions or being prepared for a spawning night.

**Figure 2.**
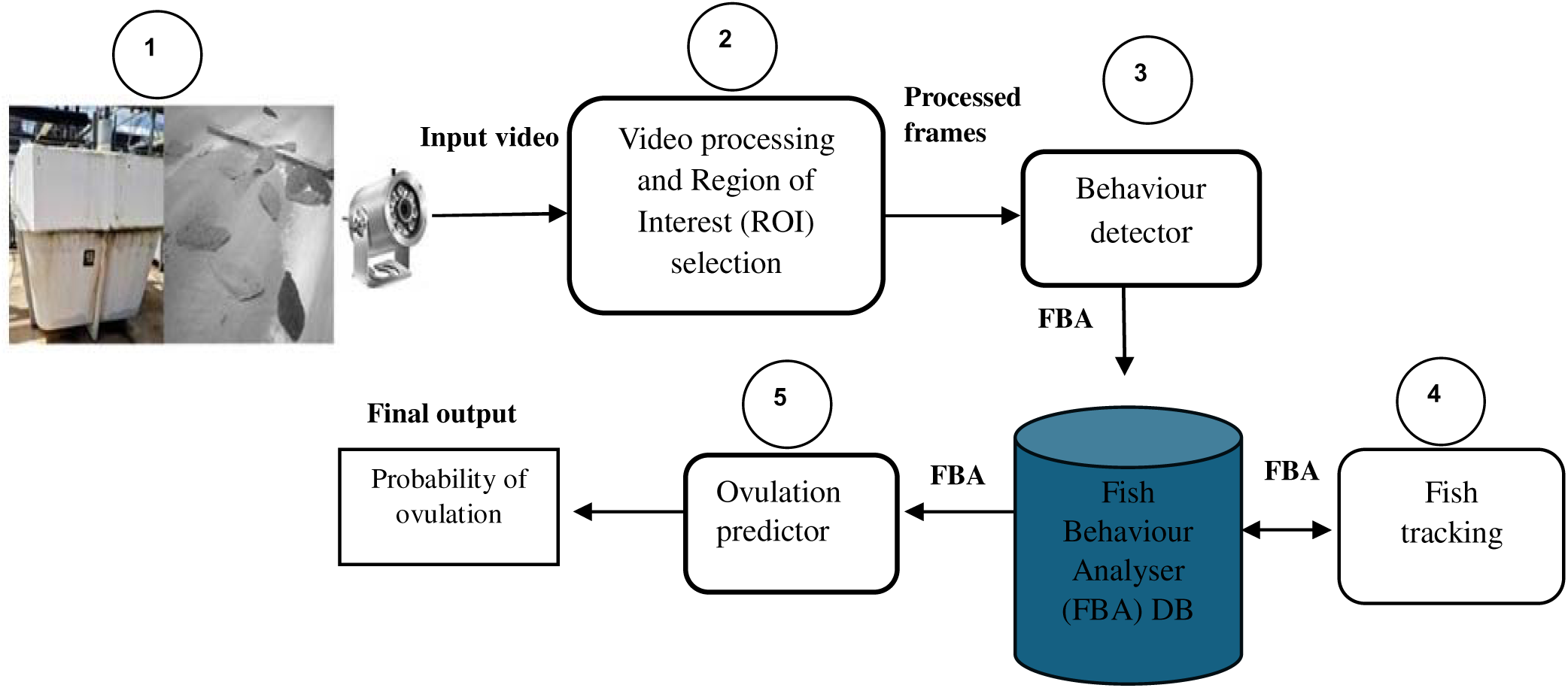
General schema diagram of the behaviour detection model, illustrating the transition from video feed capture to the prediction of the probability of ovulation.

The ovulation predictor model follows these steps:

#### Step 1. Video Input

The input of our model is the video captured by an underwater camera positioned inside the tank.

#### Step 2. Video Processing and Region of Interest (ROI) selection

The system processes the video by initially extracting each frame, then, computer vision filters are applied to the frames to enhance image quality. Finally, Regions of Interest (ROIs) are extracted from the frames. A ROI is a region of the image in which the system predicts that there is a fish. In the following steps, the system focuses only on the ROI for the behaviour analysis.

#### Step 3. Behaviour Detector

The Behaviour Detector module uses only the ROI to detect fish and assigns behavioural appearances based on their position within the frame. By focusing only on these targeted areas, this module avoids analysing the entire frame, which reduces processing time. The Behaviour Detector classifies ROIs into three classes RTH, FA and “no fish”. Only if RTH or FA categories are predicted, it generates a data structure called Fish Behaviour Analyser (FBA), which is partially filled in this module and partially filled in the next one. It consists of the following fields:

- FBA ID: A unique identifier for each behaviour entry.
- Position of Behaviour: The (x, y) coordinates where the behaviour occurs.
- Class Label: A label denoting whether the behaviour is FA or RTH.
- Timestamp: The time at which the behaviour was detected.
- Number of RTH instances: This number is initialised to zero and is updated as RTH detections accumulate over time. This field is updated directly by the Behaviour Detector.
- Number of LA instances: Initially set to zero, this value is later updated as movement-based behaviours are detected by subsequent modules. Any movement by a fish, even when not performing RTH, is counted as LA. The system logs any detected motion as an LA instance. These two metrics are independent: LA tracks all movement, while RTH tracks static periods.

The FBA database serves as a central repository for storing and retrieving all behavioural data related to fish behaviours. Each detected instance is recorded in the FBA database.

#### Step 4. Fish Tracking

The Fish Tracking module retrieves FA and RTH instances, along with position and timestamp, from the FBA database to track movement fish over time and deduce LA behaviour. This module also assigns and maintains unique Fish IDs for each detected individual, including those fish exhibiting RTH behaviour. Even in the RTH position, the module continues to track each fish individually and captures any movement they exhibit. This method filters out incidental motion caused by camera noise or minor drifting and ensures the behaviour detection remains biologically meaningful. The module updates the FBA database with the information on LA behaviour.

#### Step 5. Ovulation Predictor

The Ovulation Predictor module predicts whether fish are going to ovulate during the night using a logistic regression model. To do so, the model takes three behavioural indicators from the FBA: number of RTH behaviours, number of LA and time of the behaviour occurrence. The ovulation is predicted using the following equation:

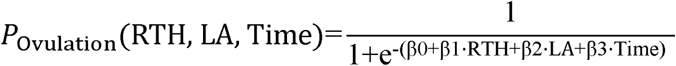

**Equation 1. Logistic regression function for ovulation prediction. P_Ovulation_ estimates the probability of ovulation based on three behavioural indicators: number of RTH events, number of LA events, and Time of observation (Time)**

Where RTH and LA represent the counts of behaviours observed. Moreover, Time is the exact time in hours at which the behaviour occurs. They are multiplied by the learned weights β₀, β₁, β₂ and β₃. The logistic function transforms the linear combination of input features into probability values. The linear equation (β₀ + β₁·RTH + β₂·LA + β₃·Time) has a domain of (-∞, +∞) since the learnable weights (β₀, β₁, β₂, β₃) can take any real values and RTH, LA, and Time are positive real numbers. The logistic transformation maps this unbounded linear output to the probability range [0, 1]. The probability P_Ovulation_ approaches 0 as the linear combination approaches -∞ and approaches 1 as it approaches +∞, though these extreme values are never exactly reached for finite input values. Values approaching 1 predict ovulation and values approaching 0 predict no ovulation. The negative exponential ensures that regardless of the magnitude of behavioural indicators, the output is always interpretable as a valid probability.

### Statistics

Sperm quality parameters were compared among male groups using one-way analysis of variance (ANOVA) with Tukey’s post-hoc tests when significant effects were detected. An independent t-test was used to compare motility between high-quality and low-quality sperm groups. Results are presented as mean ± standard deviation (SD), and statistical significance was accepted at P < 0.05.

The effects of night category (ovulation nights vs. no ovulation nights) and time (16:00–20:00 h) on RTH and LA were evaluated using two-way analysis of variance (ANOVA). The interaction between night category and time was also tested. Assumptions of normality and homogeneity of variances were verified using Shapiro–Wilk and Levene’s tests, respectively. When significant effects were detected, Tukey’s post-hoc tests were applied. Statistical significance was set at P < 0.05.

## 3. Results

This section is divided into two main parts. The first part (Sections 3.1, 3.2, 3.3) presents the proof of concept to apply IVF to naturally ovulated eggs, including spawning performance, sperm and egg quality, and behavioural patterns associated with reproductive activity. The second part (Sections 3.4, 3.5) focuses on the outcomes of the artificial intelligence–based approach, specifically the automated behavioural analysis and logistic regression model used for ovulation prediction. The results obtained in the artificial system in Sections 3.4 and 3.5 facilitate in the success of IVF procedures tested in the proof of concept in Sections 3.1, 3.2 and 3.3, with an important reduction in human effort.

### 3.1 Natural Spawning Quality

Figure 3 presents the spawning recorded throughout the experimental period in both Mix1 and Mix2 groups, distinguishing between fertilised and unfertilised spawning events. Fertilisation success was low, with only five fertilised spawns recorded across both groups during the entire study. In group Mix1, spawning was observed on only five days: two days with fertilised spawns and three with unfertilised spawns, while no spawning occurred on the remaining 47 days. The fertilised spawns were recorded on 25 and 26 March with an average fertilisation rate of 15%. On 25 March, a total egg volume of 110 mL (70 mL floating, 40 mL non-floating) was obtained at a temperature of 16.6°C, and on 26 March, 20 mL of eggs were collected (all floating) at a temperature of 16.2°C. The highest spawning volumes were recorded on 20 March with 120 mL (60 mL floating, 60 mL non-floating). Group Mix2, gave three successful fertilised spawns that were recorded on 22 March, 16 April, and 20 April with an average fertilisation rate of 10%. The spawning rate was higher in group Mix2; out of 51 monitored days, Mix2 had 24 days with spawns and 27 days with no spawns. The highest egg volumes were recorded on 6 April (230 mL; 180 mL floating, 50 mL non-floating) at a temperature of 16.7°C. Another significant volume was observed on 20 March (175 mL; 90 mL floating, 85 mL non-floating) at a temperature of 15.2°C. The last spawning event was recorded for group Mix2 on 5 May with a total egg volume of 100 mL (40 mL floating, 60 mL non-floating) at a temperature of 19°C.

**Figure 3.**
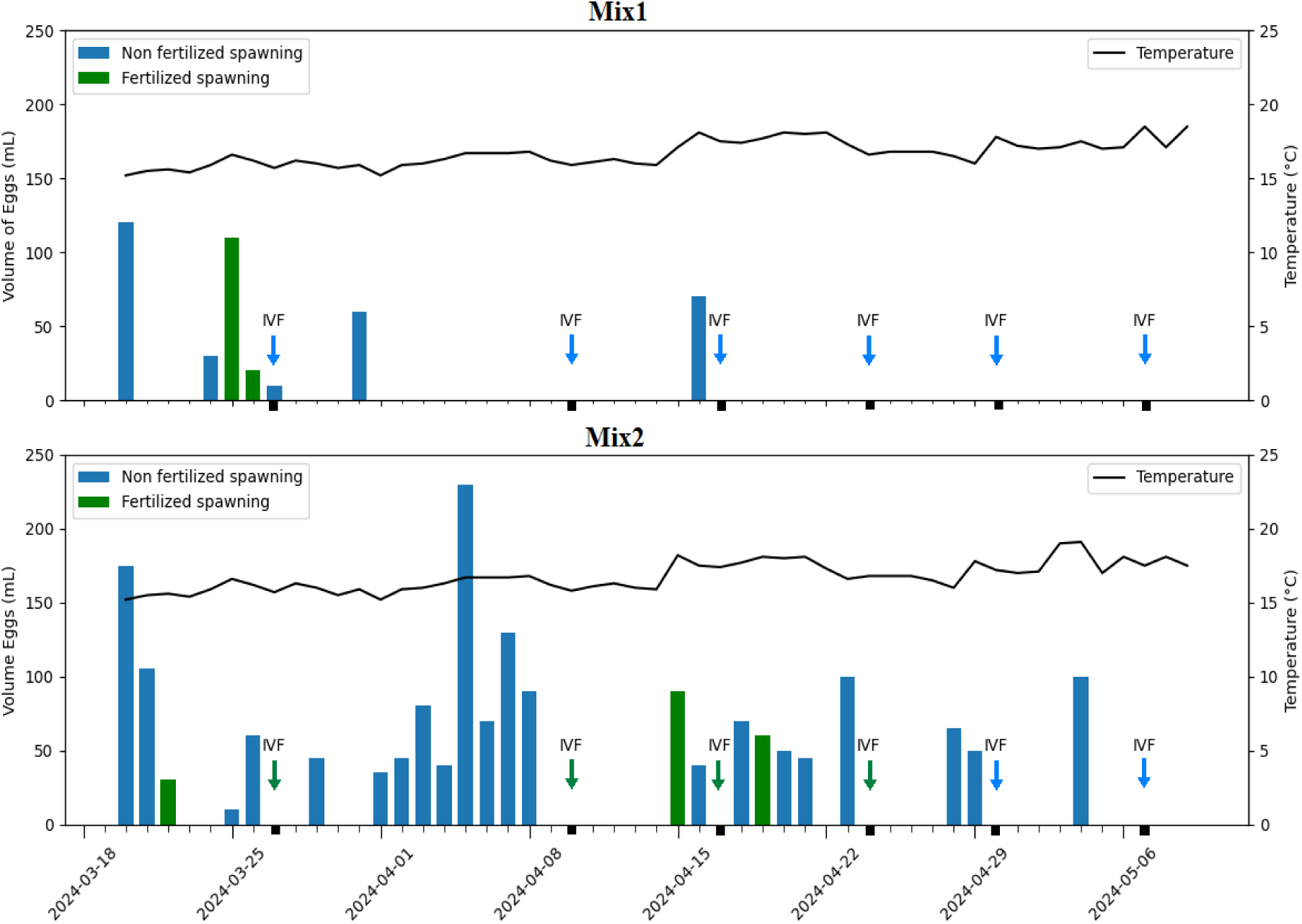
Total egg volume (mL) collected each day from Senegalese sole (*Solea senegalensis*) breeders, along with temperature data, during the 2024 spawning season for all treatments: Group Mix1 and Group Mix2. The green bars indicate fertilised eggs and the blue bars represent spawns with no fertilisation. Arrows labelled with ‘IVF’ mark the days when IVF was performed. The green arrows represent when at least one IVF treatment gave fertilised eggs, and blue arrows represent when no IVF treatments were successful.

Overall, fertilisation rates remained low in both groups throughout the study, and no fertilised spawns were observed after 20 April. Fertilised spawns confirmed that females ovulated during the night before the spawns were collected. Spawning was often observed without fertilisation, which indicated that females ovulated during the night preceding the collection of unfertilised eggs. No egg release was recorded after a significant number of nights, suggesting that ovulation did not occur on those nights.

### 3.2 Sperm collection and quality assessment for IVF

A total of 51 males were evaluated across groups Mix1 and Mix2 to assess sperm quality (Supplementary Table S1). Based on motility and velocity parameters, males were classified into three categories: Low quality sperm that was rejected for use (n = 24), High quality sperm not used for IVF (n = 12), and High quality sperm used for IVF (n = 15). A total of 24 males were excluded due to poor sperm parameters, 12 males presented high quality sperm not used for IVF, and 15 males with high quality sperm used for IVF were selected based on their superior sperm concentration, motility and velocity parameters. Significant differences were detected among the three groups for most sperm quality parameters. Males in the Low quality sperm group exhibited motility of 36.8 ± 14.5%, which was not significantly different from the High quality sperm not used for IVF group (42.9 ± 16.4%, P = 0.469). However, both groups showed significantly lower motility compared to the High quality sperm used for IVF group (69.1 ± 13.2%, P < 0.001 for both comparisons).

Similar patterns were observed for velocity parameters. The Low quality sperm group showed VCL of 47.3 ± 24.7 µm/s, VAP of 32.3 ± 21.9 µm/s, and VSL of 24.3 ± 20.8 µm/s, which were not significantly different from the High quality sperm not used for IVF group (VCL: 39.5 ± 20.6 µm/s; VAP: 25.5 ± 19.2 µm/s; VSL: 18.9 ± 17.7 µm/s; all P > 0.6). The High quality sperm used for IVF group exhibited significantly higher values for all velocity parameters (VCL: 80.1 ± 28.6 µm/s; VAP: 63.9 ± 29.9 µm/s; VSL: 55.4 ± 29.3 µm/s) compared to both other groups (all P < 0.001). Kinematic parameters including STR, LIN, WOB, and BCF followed the same pattern, with the High quality sperm used for IVF group showing significantly higher values compared to both the Low quality sperm and High quality sperm not used for IVF groups (P < 0.01 to P < 0.001), while no significant differences were observed between the Low quality sperm and High quality sperm not used for IVF groups. No significant differences were observed among groups for ALH (P = 0.132) or concentration (P = 0.055).

To evaluate short-term storage viability, the high quality sperm samples were re-analysed after 24 hours. Of the 15 males with high quality sperm, 12 had complete data for comparison. Sperm motility declined significantly from 69 ± 13.2% to 47.0 ± 17.4% (paired t-test, t = 6.746, df = 11, P < 0.001), representing a mean reduction of 21.1 ± 10.9%. Despite this decline, sperm maintained acceptable quality for fertilisation.

### 3.3 Female Ovulation Assessment and Egg Collection for IVF

During the six IVF nights, a total of 21 unique females were assessed for ovulation. Across all assessments, 16 females showed no eggs or signs of ovulation, while 5 females produced eggs as shown in Table 1. As females were checked on multiple nights, a total of 92 female assessments were conducted. Of these assessments, 34 (37%) resulted in females with eggs, while 58 (63%) showed females without eggs, releasing only fluid or showing no visible signs of ovulation. The proportion of females with eggs varied considerably across nights, ranging from 63% (10 out of 16 females) on 10 April to only 13% (2 out of 15 females) on 7 May. Total viable egg production also varied substantially, from 478.9 mL on 27 March to only 40 mL on 7 May. Successful fertilisation was achieved exclusively from group Mix2 females on four of the six IVF nights (27 March, 10 April, 17 April, and 24 April). Group Mix1 females, despite being assessed on all six nights, produced no fertilised eggs. The two nights with lowest egg availability (30 April and 7 May) yielded no fertilisation success from either group.

**Table 1.** Summary of IVF results, including the number of eggs, sperm motility, fertilisation rates, and hatching success across eight females. The table presents the total number of eggs, sperm concentration, fertilisation percentages (± standard deviation), and the resulting number of larvae, highlighting variability in reproductive success.

| Date | Tag Id | Fertilized technique | Time (h) IVF | N<sup>o</sup> eggs in 0,5 mL | Egg Volume (mL) | Total No. of Eggs obtained | N<sup>o</sup> males used | Total No. of Motile Sperm ($\times 10^6$) | No. of Sperm per Egg | % Sperm Motility ± S.D. | % Fertilized ± S.D. | Total No. Fertilized eggs | % Hatch ± S.D. | Total No. larvae eggs |
| --- | --- | --- | --- | --- | --- | --- | --- | --- | --- | --- | --- | --- | --- | --- |
| 27/3/24 | F834 | 100 mL beaker | 23:59 | 264 | 170 | 89760 | 3 | 4M | 15152 | 72,7 ± 15,2 | 38 ± 14 | 35006 | | |
| 27/3/24 | 1B4A | 100 mL beaker | 0:15 | 305 | 100 | 61000 | 3 | 4M | 13115 | 72,7 ± 15,2 | 34 ± 19 | 20740 | | |
| 27/3/24 | 3193 | 100 mL beaker | 0:45 | 144 | 41.5 | 11952 | 2 | 4M | 27778 | 77 ± 13,3 | 23 ± 6 | 2749 | | |
| 27/3/24 | 82C1 | 100 mL beaker | 0:57 | 156 | 44.2 | 13790.4 | 2 | 4M | 25641 | 77 ± 13,3 | 8 ± 9 | 1096 | | |
| 27/3/24 | F834 | Massive | 23:59 | 264 | 170 | 89760 | Pool-2 | 80M | 303030 | 72,0 ± 21,9 | 44.4 ± 37.2 | 960 | 5.1% | 4578 |
| 27/3/24 | 1B4A | Massive | 0:15 | 305 | 100 | 61000 | Pool-2 | 40M | 131148 | 72,0 ± 21,9 | 21.97 ± 35.3 | 653 | 2.20% | 1342 |
| 10/4/24 | F834 | 100 mL beaker | 22:38 | 149 | 20 | 5960 | 3 | 4M | 26846 | 53.9 ± 1.9 | 15 ± 10 | 64 | | |
| 10/4/24 | F834 | Massive | 22:38 | 149 | 20 | 5960 | Pool-2 | 80M | 536913 | 53.9 ± 1.9 | 4.6 ± 1.6 | 64 | 7% | 417 |
| 17/4/24 | 8486 | 100 mL beaker | 23:40 | 153 | 21 | 6426 | 2 | 4M | 26144 | 84.5 ± 5.2 | 1 ± 2 | 69 | | |
| 17/4/24 | 8486 | Massive | 23:40 | 153 | 100 | 30600 | 1 | 80M | 522876 | 84.5 ± 5.2 | 0.01 ± 0.0043 | 327 | 0.70% | 214 |
| 24/4/24 | F834 | 100 mL beaker | 23:32 | 245 | 81.5 | 39935 | 1 | 4M | 16327 | 72 ± 3.6 | 6 ± 8 | 427 | | |
| 24/4/24 | F834 | Massive | 23:32 | 245 | 19.5 | 9555 | 1 | 80M | 326531 | 72 ± 3.6 | 21.7 ± 3 | 102 | 18% | 1720 |

From the females with eggs, 70 female–male pairing instances were created for fertilisation attempts. Of these, 21 instances resulted in successful fertilisation across four nights, involving five females, while 49 paired crosses showed no fertilisation. The first IVF night (27 March 2024) was the most successful, with eggs fertilised from four females (four beaker fertilisations and two massive fertilisations), while eggs from four females remained unfertilised. On the subsequent three successful nights, one female per night produced fertilised eggs, with fertilsation using the two methods (beaker and massive fertilisation per female). The remaining two IVF nights (30 April and 7 May) produced no successful fertilisation despite multiple female–male pairings being attempted.

Table 1 presents IVF outcomes using two methods: beaker fertilisation (individual male-female pairings) and massive fertilisation (pooled sperm from multiple males). A total of 26 unique males were used, with eight contributing to successful fertilisation. Some males were reused on different days, resulting in 51 male usage instances, of which 21 resulted in successful fertilisation. In beaker fertilisations using 4 million motile sperm per trial, fertilisation rates varied significantly across females, ranging from 38% in Female F834 (35,006 fertilised eggs from 89,760) to 1% in Female 8486 (69 fertilised eggs from 6,426), despite the latter having the highest sperm motility (84.5%). In massive fertilisation trials, where pooled sperm was used, fertilisation rates ranged from 0.01% to 44.4%, with the highest success in Female F834 producing 4,578 larvae (5.1% hatching rate). Female F834 also achieved the highest hatching percentage (18%) on 24 April with a fertilisation rate of 21.7%, yielding 1,720 larvae.

The difference in IVF success between groups reflected their natural spawning frequencies. Group Mix2, which had higher natural spawning activity (24 spawning days out of 51 monitored, 47%), produced all fertilised eggs during IVF trials. Group Mix1, with substantially lower spawning frequency (5 spawning days out of 52 monitored, 10%), produced no fertilised eggs despite females being assessed on all six IVF nights. Since IVF nights were selected without prior knowledge of ovulation status, success depended on whether females happened to have ovulated. The higher ovulation frequency and spawning in Mix2 increased the probability of encountering ovulated females on randomly selected nights, whilst the low ovulation frequency in Mix1 resulted in repeated missed opportunities.

### 3.4 Automated Behaviour Analysis during peak Reproductive Hours

The average hourly RTH counts (16:00–20:00 h) differed significantly among the five-night categories—ovulation with fertilised eggs, IVF-fertilised nights (only tank Mix2), ovulation with unfertilised eggs, no-ovulation nights, and IVF-unfertilised nights—in both experimental groups (Mix1: F(3,70) = 3.079, P = 0.033; Mix2: F(3,85) = 2.936, P = 0.038) (Figure 4). RTH activity also varied significantly across time in both groups (Mix1: F(4,70) = 3.860, P = 0.007; Mix2: F(4,85) = 4.389, P = 0.003), with the highest mean values occurring between 18:00 and 19:00 h. Although ovulation nights generally showed higher hourly RTH activity than no-ovulation nights, there was overlap in standard deviations and no significant interaction between night category and time was detected (Mix1: P = 0.982; Mix2: P = 0.997), indicating similar temporal patterns across all four-night types.

**Figure 4.**
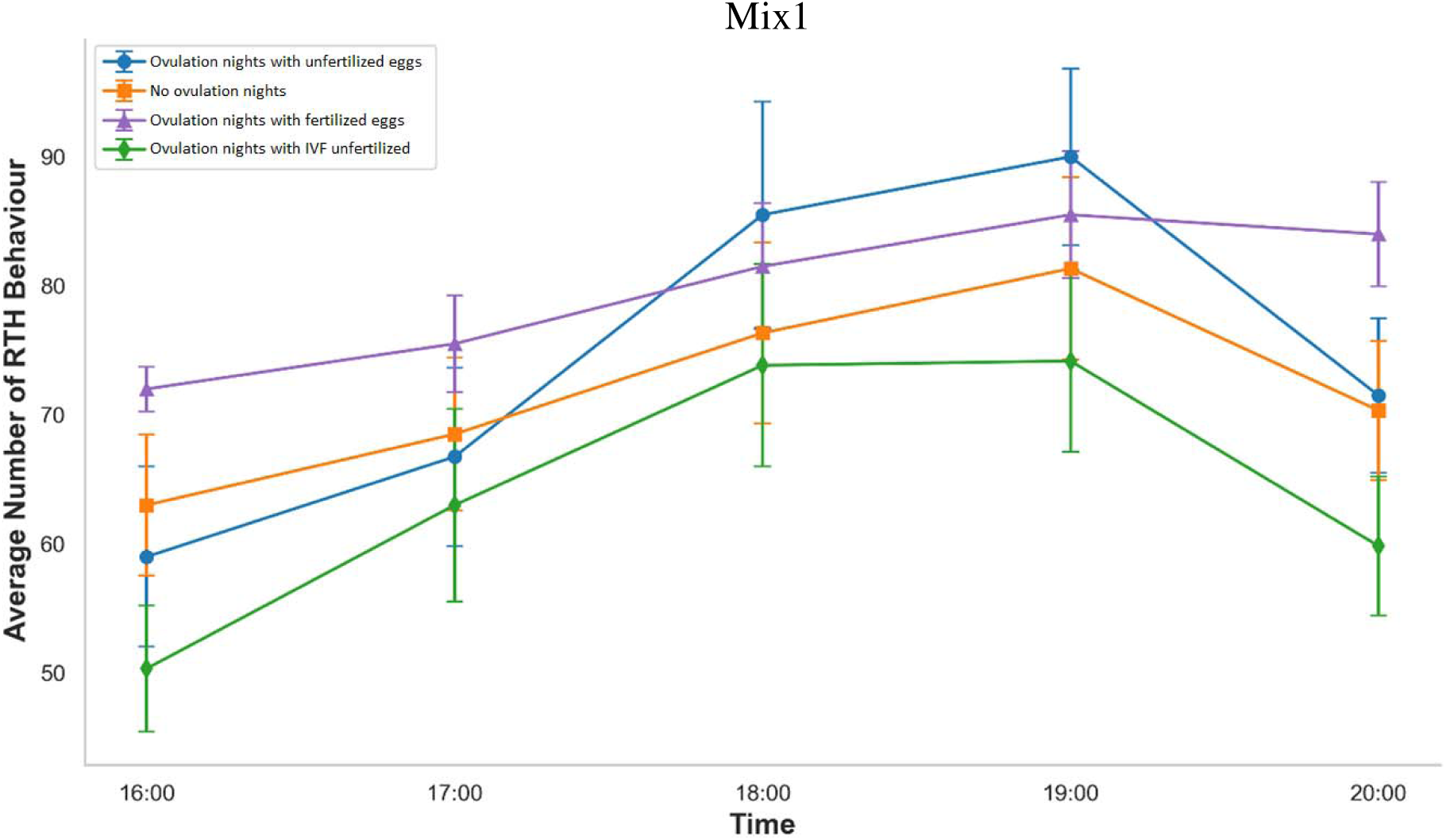
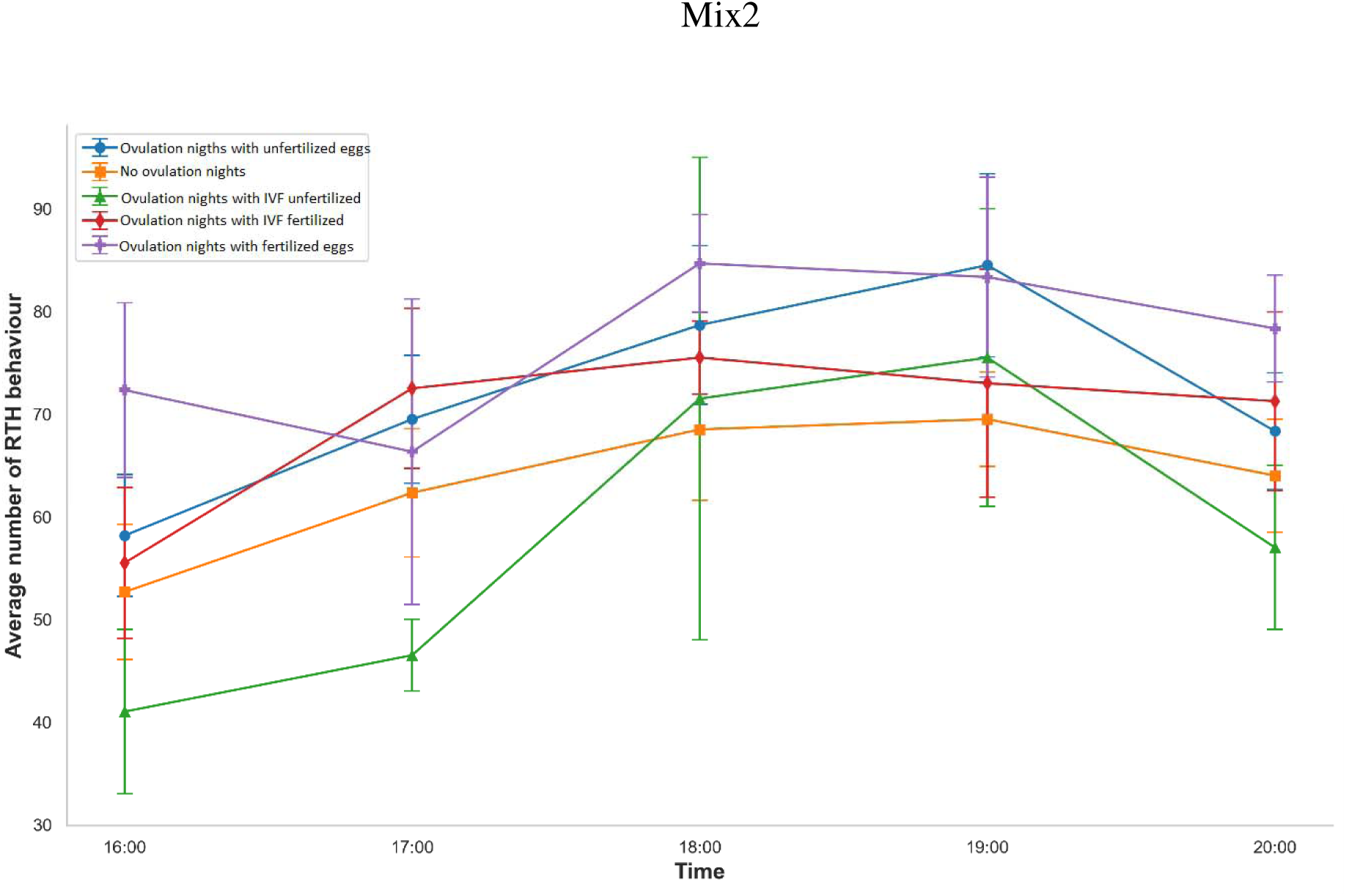
Average number of RTH behaviour occurrences over six spawning (fertilised, unfertilised) and six non-spawning nights for group Mix1 (above) and Mix2 (below), recorded between 16:00 and 20:00. A two-way ANOVA revealed significant effects of both night type and time on RTH behaviour (P < 0.05 for both), but no significant interaction between night type and time.

Ovulation nights with fertilised eggs exhibited high RTH activity, reaching 85.5 ± 4.91 at 19:00 h in Mix1 and 84.67 ± 4.75 at 18:00 h in Mix2. Ovulation nights with unfertilised eggs showed a numerically higher peak at 19:00 h (90.5 ± 7.00) before decreasing to 71.5 ± 6.2 by 20:00 h. Tukey’s post-hoc test confirmed that ovulation nights with fertilised eggs had significantly higher RTH values than ovulation nights with IVF unfertilised eggs (mean difference = 15.467, P = 0.049). RTH activity was significantly higher at 19:00 h (LS Mean = 81.67) and 18:00 h (LS Mean = 78.71) compared to 16:00 h (LS Mean = 61.08) (P = 0.007 and P = 0.031, respectively).

Although not statistically significant, ovulation nights with fertilised eggs had a higher mean RTH (77.0 ± 4.29) than no ovulation nights (63.4 ± 3.04; P = 0.055). Ovulation nights with unfertilised eggs also displayed elevated activity, peaking at 84.5 ± 8.91 at 19:00 h. The IVF-related categories, including ovulation nights with IVF fertilised eggs (Mix2) and IVF unfertilised, showed comparable peaks at 75.5, though the latter had greater variability (±14.5). This temporal trend was consistent across both experimental groups, with RTH activity peaking during the late afternoon and early evening.

Overall, both Mix1 and Mix2 demonstrated a consistent pattern of elevated RTH activity during the late afternoon and early evening in association with ovulation, supporting the use of this behaviour as a non-invasive biomarker for reproductive readiness.

LA showed a stronger association with ovulation than Rest the Head (RTH) behaviour in both groups (Mix1: F(3,70) = 6.388, P < 0.001; Mix2: F(3,85) = 18.054, P < 0.001) (Figure 5). The overall average LA was highest on ovulation nights, with peak activity generally recorded between 19:00 h and 20:00 h.

**Figure 5.**
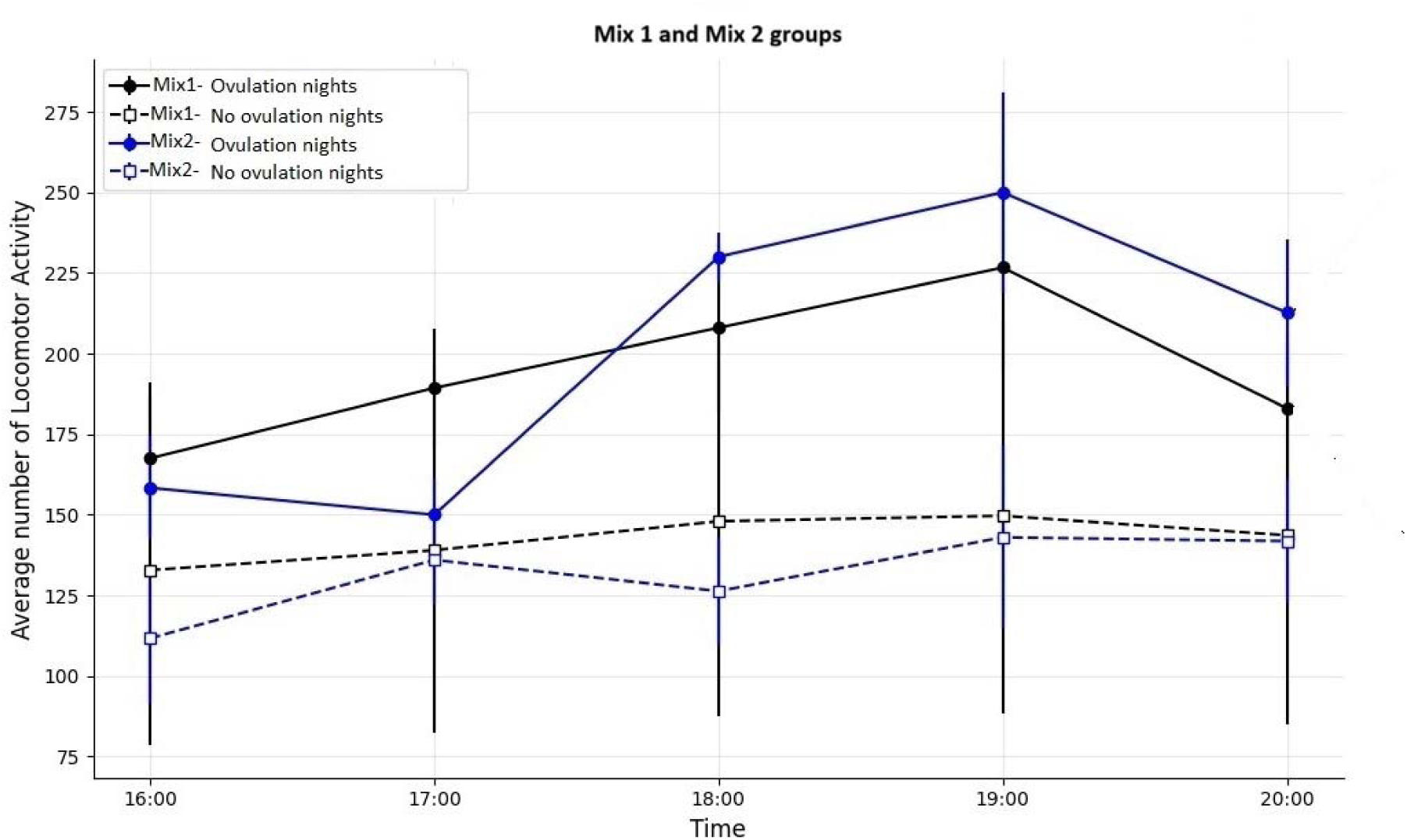
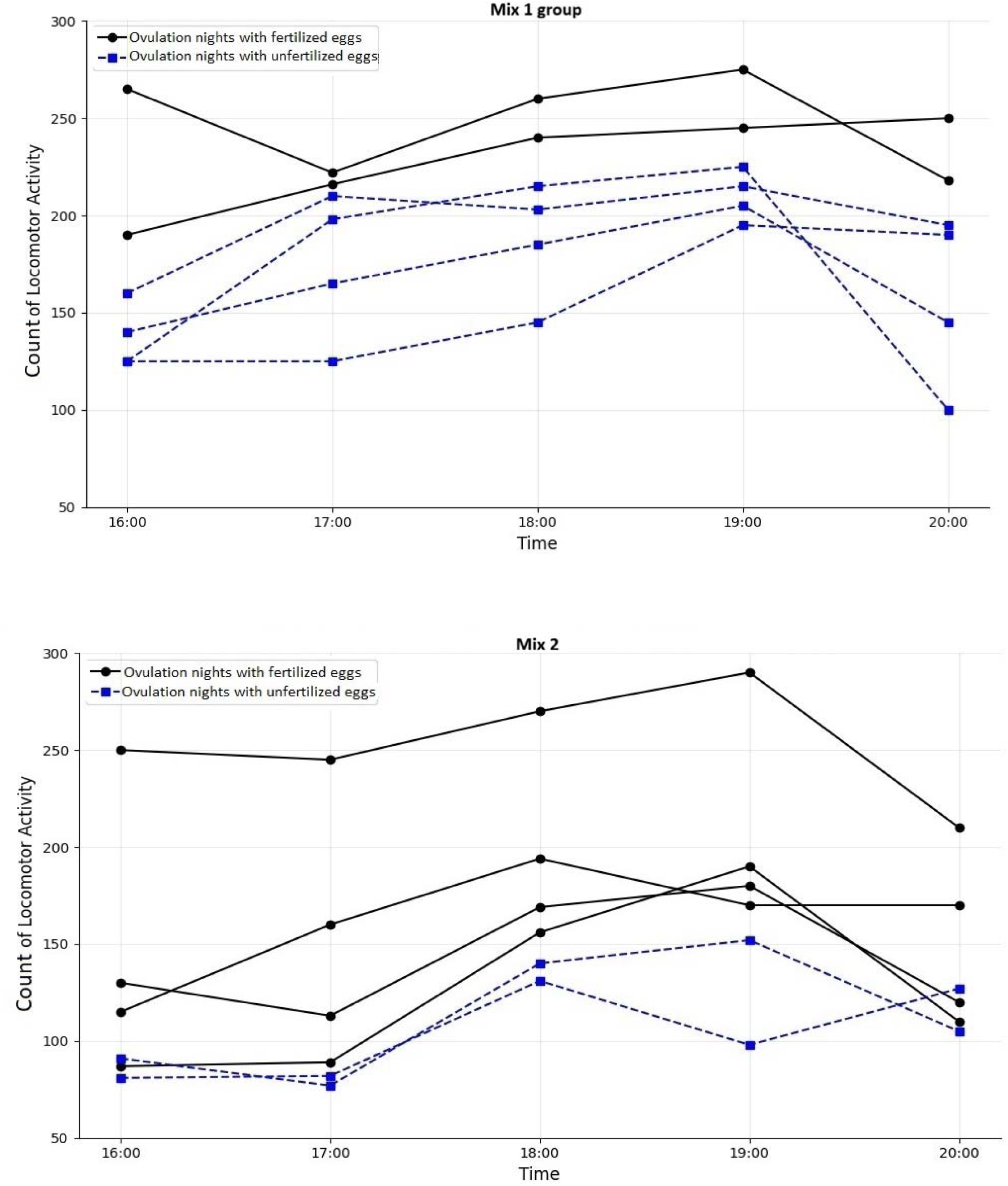
LA patterns in Senegalese sole during ovulation and non-ovulation nights. (above) Average number of movements per fish and hour (16:00–20:00 h) for all ovulation nights (n = 6) and all no ovulation nights (n = 6) in groups Mix1 and Mix2. (middle) Total movement counts per hour (16:00–20:00 h) for individual ovulation nights, nights with fertilised eggs (n = 2) versus unfertilised eggs (n = 4) in group Mix1. (below) Total movement counts per hour (16:00–20:00 h) for individual ovulation nights, nights with fertilised eggs (n = 4) versus unfertilised eggs (n = 2) in group Mix2.

LA increased on ovulation nights in Mix1 group, with the highest values recorded between 19:00 h and 20:00 h (Figure 5 - middle). The least squares mean (LS Mean) LA peaked at 260.00 movements at 19:00 h on Spawn Fertilised nights, whereas No Spawn nights showed lower activity levels, with the highest LS Mean reaching only 149.67 movements at the same time. In addition, nights with fertilised eggs showed higher LA, with activity peaking at 290 movements, compared with unfertilised nights, which ranged from 98 to 152 movements.

Finally, LA increased progressively from 16:00 h onwards in Mix2, peaking at 269.67 movements at 19:00 h on Spawn Fertilised nights and 250.00 movements at the same time on Spawn Unfertilised nights (Figure 5 - lower). In contrast, on no-ovulation nights, LA remained lower throughout the observation period, ranging from 111.67 movements at 16:00 h to 143.00 movements at 19:00 h. As in Mix1, nights with fertilised eggs showed higher LA than unfertilised nights, with peak values ranging from 275 to 292 movements, while unfertilised nights showed lower peaks between 195 and 215 movements.

Overall, these results indicate that fertilised ovulation nights coincide with the general LA peak observed around 19:00 h and are associated with higher activity levels in both experimental groups.

### 3.5 Performance of the ovulation prediction model

Figure 6 shows on the left the ROC curve and on the right the confusion matrix returned by the ovulation predictor applied to the combined dataset from Mix1 and Mix2. A probability threshold of 0.5 was used, that is, afternoons that the prediction was larger than 0.5 were classified as ovulation nights.

**Figure 6.**
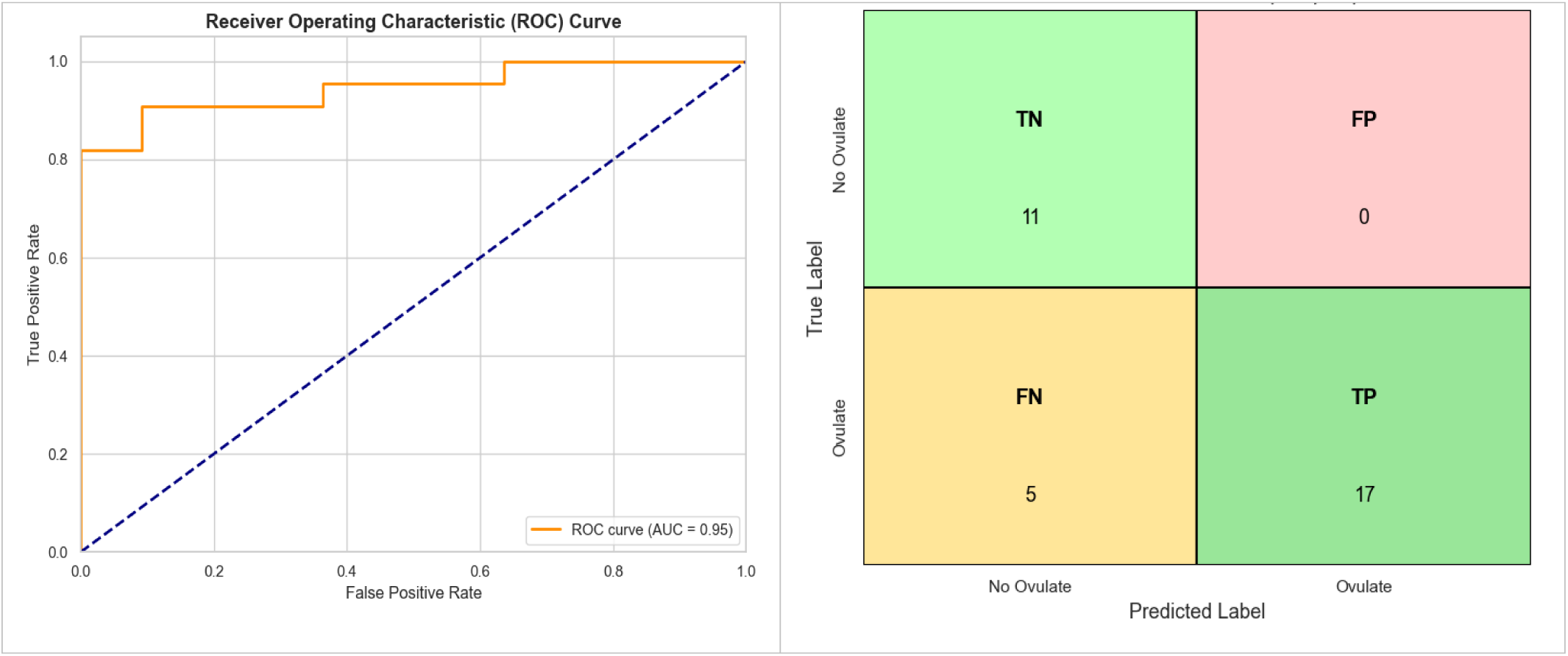
Performance of the logistic regression for ovulation classification trained on behavioural data from group Mix1 and Mix2. The probability threshold to classify ovulation nights or non-ovulation nights was set to 0.5. (left) ROC curve. (right) Confusion matrix.

Using this threshold, the ROC achieves an AUC of 0.95. Moreover, the model achieved a precision of 1.00 and recall of 0.77 for the ovulation class, yielding an F1-score of 0.87. For the nights with no ovulation, the model achieved a recall of 1.00 and precision of 0.69, indicating perfect detection of non-ovulation nights but with some false positives.

The model correctly identified 17 out of 22 ovulation nights and all 11 non-ovulation nights, achieving an overall accuracy of 85%. This confirms the model’s strong discriminative ability, particularly its high recall for both classes, ensuring that ovulation events are rarely missed while all non-ovulation nights are correctly identified.

Following the evaluation on the combined dataset, the same trained model was evaluated separately on each experimental group to assess group-specific performance. Using the behavioural inputs RTH, LA, and Time, a combination of probability thresholds was systematically tested for each group to identify the optimal threshold that maximized classification accuracy.

Figure 7 presents the group-specific performance of the logistic regression model for Mix1 and Mix2, showing the classification results obtained using the optimal probability threshold for each experimental group. For group Mix1, among the different thresholds evaluated, 0.5 showed the best performance. Out of 20 unique nights, the model correctly predicted all 9 nights with ovulation and 6 out of 11 no ovulation nights, resulting in a prediction accuracy of 75%. The classification report further confirmed this outcome, showing a perfect recall of 1.00 for ovulation nights and a precision of 1.00 for no ovulation nights, indicating the model’s ability to detect all spawning events whilst minimising false positives (Figure 7a). For group Mix2, evaluation of different thresholds determined that 0.28 was the optimal threshold for this group. Using this threshold, out of 17 nights analysed, the model correctly predicted 8 out of 9 ovulation nights and all 8 no ovulation nights, achieving an overall accuracy of 82%. The classification report showed a precision of 1.00 for ovulation nights and a perfect recall of 1.00 for no ovulation nights, highlighting the model’s reliability in identifying nights with no ovulation (Figure 7b). Overall, the regression model performed well across both experimental groups, with group Mix2 achieving higher accuracy than group Mix1 (82% vs. 75%), demonstrating the robustness of the approach across different group conditions.

**Figure 7.**
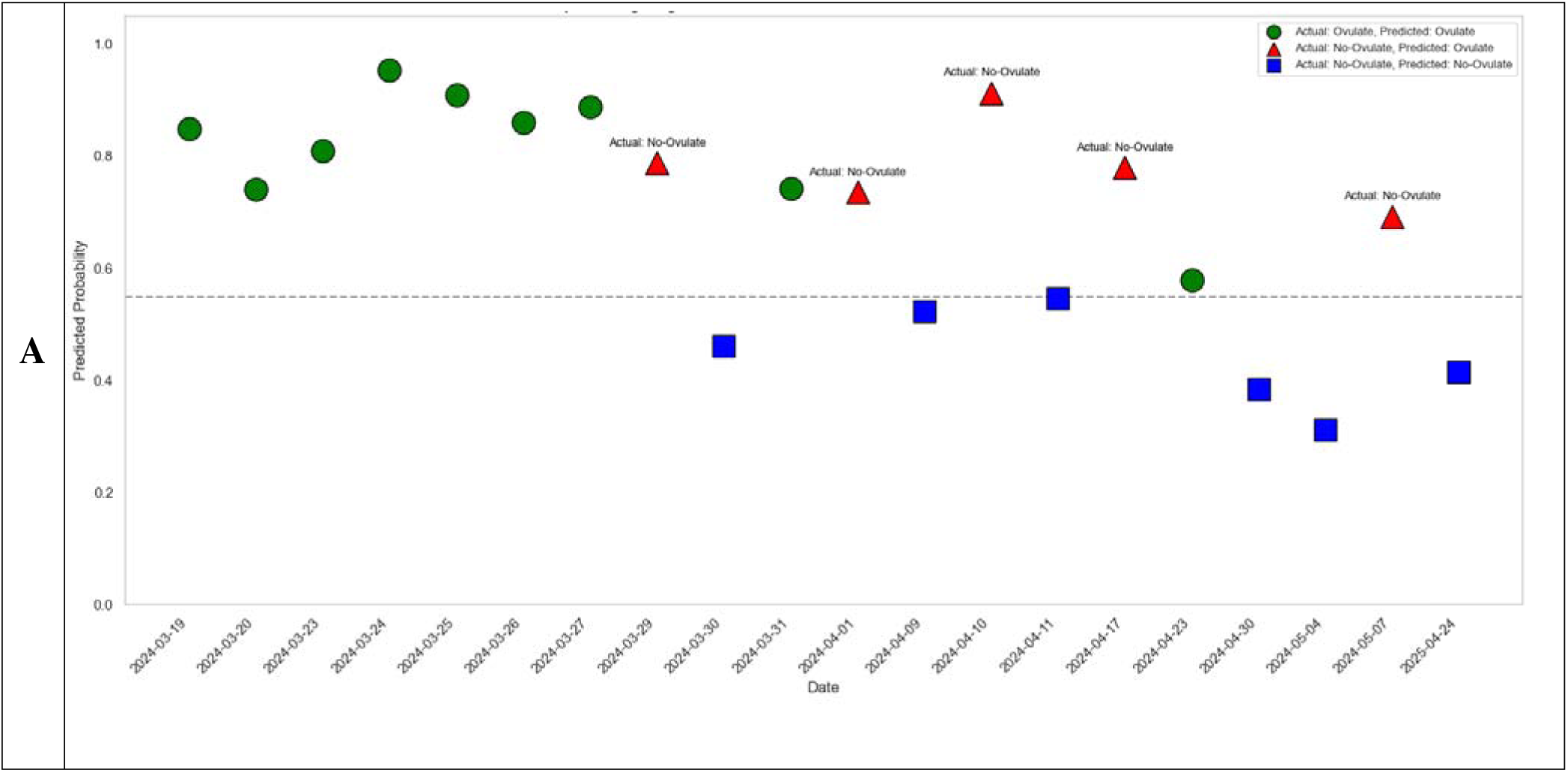
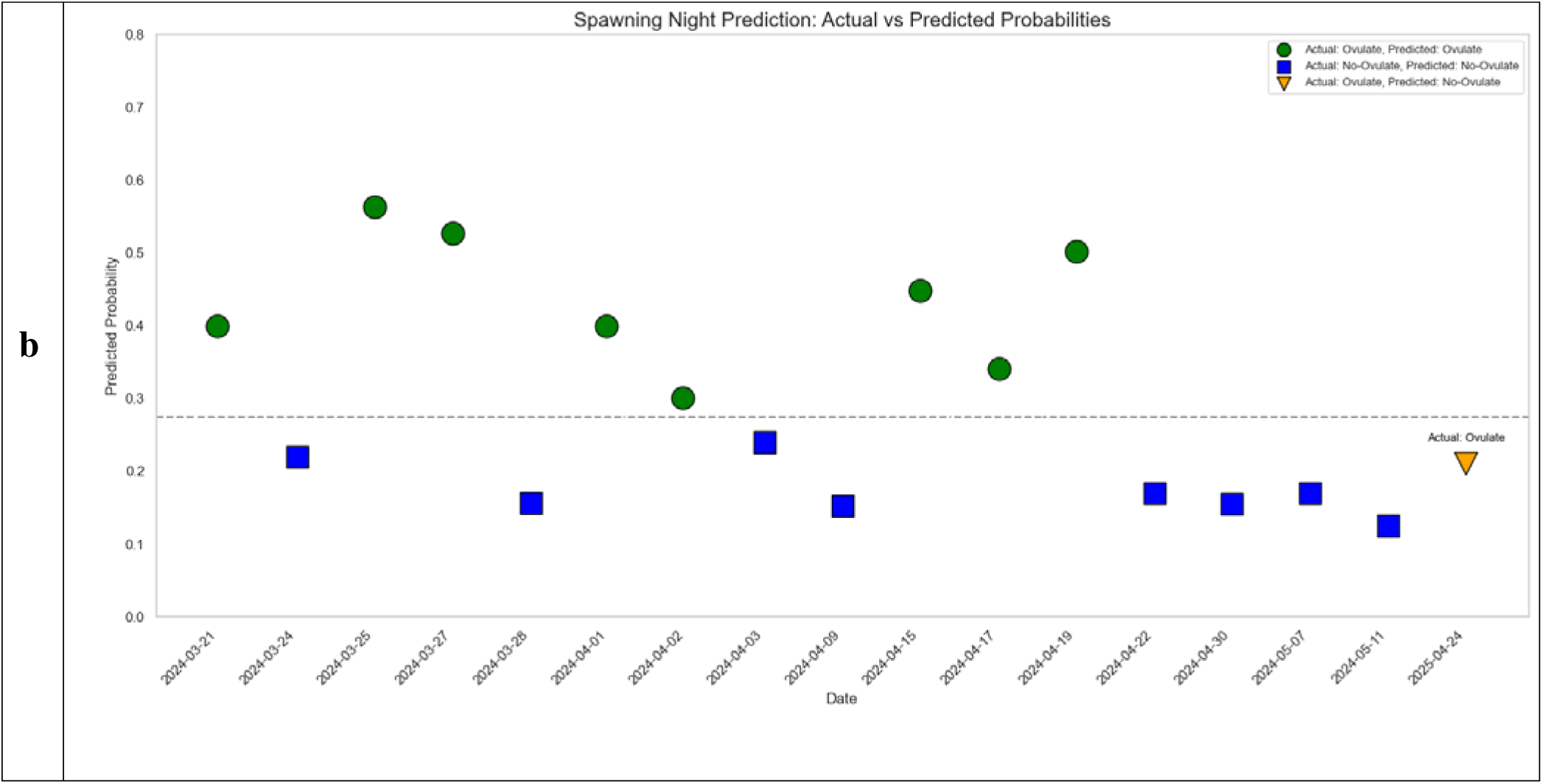
Predicted probabilities of ovulation and non-ovulation nights based on logistic regression model. Each point represents the model’s predicted probability of ovulation for a given night, showing both actual and predicted classification outcomes. Green circles and blue squares indicate correct predictions for ovulation nights and no ovulation nights, respectively. Orange downward triangles represent misclassified ovulation nights, and red upward triangles indicate misclassified no ovulation nights. A dashed horizontal line marks the decision threshold separating predicted ovulation (above) from predicted non-ovulation (below). (a) Group Mix1 across 20 nights. (b) Group Mix2 across 17 nights.

Figure 8 summarises the confusion matrix outcomes of the logistic regression model for Mix1 and Mix2. The model achieved 75% accuracy for Mix1 (TP = 9, FP = 5, FN = 0, TN = 6) and 82% accuracy for Mix2 (TP = 8, FP = 0, FN = 1, TN = 8), confirming consistent performance across both groups.

**Figure 8.**
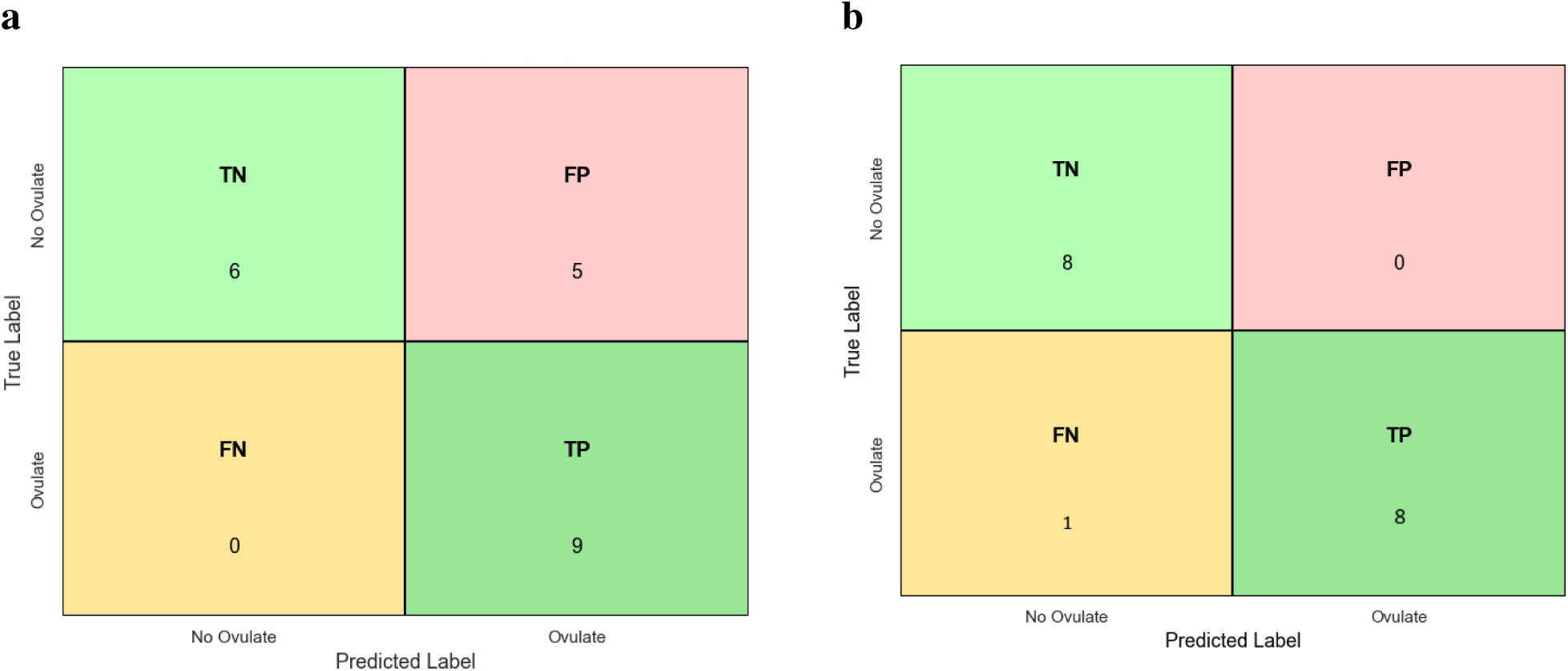
Confusion matrix evaluating the performance of the logistic regression model in predicting ovulation and no ovulation nights for the two experimental groups, given the probability threshold of 0.28. *(a)* group Mix1: the model achieved 75% accuracy, with True Positives (TP) = 9, False Positives (FP) = 5, False Negatives (FN) = 0, and True Negatives (TN) = 6. *(b)* group Mix2: the model achieved 82% accuracy, with TP = 8, FP = 0, FN = 1, and TN = 8. The matrices illustrate the distribution of true and predicted labels based on behavioural features including RTH, LA, and Time.

## 4. Discussion

This study demonstrates that hormone-free fertilisation is technically viable for Senegalese sole but requires automated behavioural monitoring to overcome the practical challenge of unpredictable ovulation timing. By combining hormone-free IVF protocols with machine learning-based ovulation prediction, this work establishes a framework for sustainable, organic aquaculture that eliminates dependence on hormonal treatments while ensuring commercial feasibility. Despite observable reproductive behaviours and spawning events in captive Senegalese sole, fertilisation success remained limited, reflecting persistent challenges in achieving natural reproduction under controlled conditions. Our findings support earlier reports that reproductive failure in cultured populations is often associated with disrupted courtship behaviour and altered social dynamics (Fatsini et al., 2020; González-López et al., 2024). These constraints, including limited parental participation and potential impacts from social hierarchies or mate choice, appear to reduce the effective breeding population and highlight the need for improved broodstock management strategies (Porta et al., 2006; Salas-Leiton et al., 2010; Martín et al., 2014). The development of hormone-free protocols is particularly relevant given the increasing demand for organic aquaculture products and regulatory restrictions on hormonal treatments in organic certification systems (European Commission, 2021a). To our knowledge, this study provides the first demonstration of successful IVF using naturally ovulated eggs without hormonal intervention in Senegalese sole, producing viable larvae with fertilisation rates up to 44% and hatching rates of 18%.

While our hormone-free fertilization experiments successfully produced viable offspring, the practical implementation revealed a critical temporal constraint that determines whether this approach can be adopted commercially. Ovulated Senegalese sole eggs maintain fertilization competence for approximately 3 hours (Rasines et al., 2012; Ramos-Júdez et al., 2021), creating a narrow window that must be precisely identified for successful gamete collection and fertilization.

In commercial aquaculture settings, manual behavioural observation would be insufficient for reliable detection of natural ovulation events. Aquaculture managers cannot feasibly maintain continuous monitoring of fish behaviour throughout evening hours during the entire spawning season, which extends for several months. The current practice of periodic evening checks is inefficient and frequently results in missed reproductive opportunities. Even with dedicated monitoring, determining the exact timing of ovulation without external indicators requires either frequent handling (which causes stress and may disrupt reproductive behaviour) or reliance on chance observation during the narrow viable window.

In the present study, IVF trials were conducted on six nights selected without prior knowledge of ovulation status, essentially representing random sampling during the spawning season. The contrasting outcomes between groups demonstrated the limitations of this approach. Females from both groups were assessed on the same six nights, yet fertilised eggs were obtained exclusively from group Mix2, which had higher natural spawning frequency (24 out of 51 days, 47%), with success on 4 of 6 nights. Group Mix1, with lower spawning frequency (5 out of 52 days, 10%), produced no fertilised eggs on any night despite repeated assessment. This disparity reflects the underlying difference in ovulation frequency: when nights were selected randomly, the probability of encountering ovulated females was inherently higher in the group where females ovulated more frequently. These results demonstrate that random selection of IVF nights is unreliable and inefficient for commercial implementation. Both natural spawning and hormone-free IVF achieved low overall fertilisation success, though for different reasons. Natural spawning produced fertilised eggs on only 5 occasions across both groups, with fertilisation rates of 10-15% when successful, primarily limited by male behavioural dysfunction where males failed to participate in spawning. IVF trials achieved fertilisation rates ranging from 1% to 44% when ovulated eggs were present, but frequently failed entirely when females had not ovulated on the selected nights or eggs obtained were not viable as the 3h window of viability had passed. The common factor underlying both approaches was the unpredictability of reproductive events: for natural spawning, the unpredictability of male participation; for IVF, the unpredictability of female ovulation timing. Accurate prediction of ovulation would address the timing constraint for IVF, enabling gamete collection when eggs are at optimal viability.

Therefore, our automated behavioural monitoring system, which achieved 82–85% accuracy in predicting ovulation nights across both experimental groups, is not merely a technical enhancement but an essential enabling technology for making hormone-free fertilisation practically viable in commercial settings. In practical terms, this accuracy means that instead of random sampling where success depends entirely on ovulation frequency (10% in Mix1, 47% in Mix2), targeted IVF attempts guided by the prediction model could achieve success on over 80% of selected nights. For a facility conducting IVF trials, this represents a substantial improvement in operational efficiency, reducing wasted resources and increasing the proportion of nights that yield fertilised eggs. The system addresses the timing challenge by predicting ovulation several hours before it occurs, enabling aquaculture managers to prepare equipment, collect fresh sperm from selected males, and monitor specific females for optimal gamete collection. This transforms the process from an unpredictable approach reliant on chance observation to a managed protocol where resources can be strategically allocated.

The practical implications are significant for organic aquaculture certification. Without reliable ovulation prediction, hormone-free fertilization remains unsuitable for commercial operations due to its unpredictability and low success rate. The integration of behavioural monitoring and machine learning provides a non-invasive solution that maintains the ethical and regulatory requirements of organic certification while delivering the reliability necessary for commercial implementation. By enabling predictive management rather than reactive response, this approach makes organic certification achievable for Senegalese sole aquaculture operations. Hormonal induction remains a widely used method to overcome reproductive dysfunction in captive Senegalese sole. Rasines et al. (2012) demonstrated that both GnRHa injections and implants effectively induce ovulation, facilitating spawning in captive broodstock. However, hormonal treatments can involve handling stress, regulatory constraints, and potential impacts on fish welfare. Our IVF experiments demonstrated that targeted male selection based on sperm quality is a crucial strategy to enhance fertilisation success. Importantly, the fertilisation failures observed in this study cannot be attributed to insufficient sperm. Each fertilisation attempt used 4 million motile spermatozoa per beaker for approximately 150-300 eggs, providing 13,000-28,000 sperm per egg. This represents a 7-17 fold excess compared to the 1,600-2,000 sperm per egg reported as sufficient for near-complete fertilisation (∼99%) in hormone-induced protocols (Ramos-Júdez et al., 2021). Furthermore, sperm quality remained acceptable after 24 hours storage (47.0 ± 17.4% motility), confirming sperm viability was maintained throughout the fertilisation process. These findings conclusively demonstrate that sperm quality and quantity were not limiting factors, and that the variable fertilisation success (0.01% to 44%) was primarily determined by female factors, including egg quality and timing of collection relative to ovulation.

These findings highlight the need for accurate prediction of ovulation timing to optimise hormone-free fertilisation protocols. Previous studies on Senegalese sole sperm have demonstrated that sperm quality varies considerably among individuals and is influenced by factors including age, diet, and season (Cabrita et al., 2006; García-López et al., 2006). The use of CASA technology for objective sperm quality assessment, as employed in this study, provides a reliable and standardised method for male selection that has been validated in multiple teleost species (Rurangwa et al., 2004). Short-term cold storage of Senegalese sole sperm has been shown to maintain fertilisation capacity for 24-48 hours when diluted in appropriate extenders (González-López et al., 2020), and which is confirmed in the present study.

Nonetheless, hormone-free IVF in our study successfully produced viable offspring. Under standard IVF conditions (100 mL beaker trials), fertilisation rates reached up to 38%, with the highest outcome observed in Female F834, which produced over 35,000 fertilised eggs. In massive fertilisation trials, fertilisation percentages were generally lower but still resulted in meaningful larval production: 4,578 larvae from Female F834 (44.4% fertilisation; 5.1% hatching) and 1,720 larvae from Female F834 on 24 April (21.7% fertilisation; 18% hatching). These fertilisation rates, whilst lower than those achieved with hormone-induced ovulation (Rasines et al., 2012; Ramos-Júdez et al., 2021), demonstrate that hormone-free IVF can produce commercially meaningful numbers of larvae. The variability in fertilisation success among females is consistent with previous reports that egg quality in Senegalese sole is highly variable and influenced by factors including female age, nutritional status, and timing of ovulation (Cabrita et al., 2006; Beirão et al., 2009). These results confirm the technical feasibility of hormone-free IVF in Senegalese sole, provided that sperm parameters and male selection are carefully optimised, and egg collection is timed appropriately relative to ovulation. These limitations reinforce the need for alternative strategies that enable hormone-free IVF and non-invasive monitoring of female gamete maturation and ovulation. In this context, the behavioural monitoring and automated prediction model developed in our study offers a promising solution by identifying ovulatory patterns based on natural reproductive behaviours, without the use of hormonal interventions. By combining a convolutional neural network (CNN) to detect specific reproductive behaviours RTH and LA with a logistic regression classifier, the model effectively predicted ovulation events based on behavioural patterns observed during the late afternoon hours. The system achieved an overall classification accuracy of 85% and an AUC of 0.95, with high precision and recall values across both experimental groups (Mix1 and Mix2). This behaviour-driven, non-invasive system allows for continuous monitoring without the need for hormonal intervention or manual handling. As such, the automated model offers a powerful tool for improving the precision and sustainability of reproductive management, enabling hormone-free ovulation monitoring aligned with organic farming principles.

The behavioural analysis further supports the use of non-invasive monitoring as a tool to assess reproductive readiness. The logistic regression analysis provided additional insights into the relative importance of different behavioural indicators. Whilst both RTH and LA were included as predictor variables, LA showed a significant positive relationship with ovulation probability, whereas RTH exhibited a weak, non-significant negative association. This suggests that LA is a more direct indicator of imminent ovulation, whilst RTH behaviour may reflect broader courtship dynamics that are less tightly coupled to ovulation timing. These differential associations demonstrate the model’s ability to distinguish between general reproductive behaviour and specific ovulation-related activity patterns, reinforcing its value as a targeted prediction tool.

The association between increased LA and reproductive events is consistent with previous observations in Senegalese sole, where swimming activity and courtship behaviours intensify during the hours preceding spawning (Carazo et al., 2016; Duncan et al., 2019; Fatsini et al., 2020). The RTH behaviour, whilst characteristic of pre-spawning courtship interactions in this species (Martín et al., 2014), may occur throughout the reproductive season regardless of imminent ovulation, which would explain its weaker predictive value compared to LA in our model. These findings build upon our previous work demonstrating that machine learning analysis of RTH and LA behaviours can successfully predict spawning nights in Senegalese sole with high accuracy (Qadir et al., 2025). The present study extends this approach by applying the prediction model specifically to hormone-free IVF protocols, demonstrating its practical utility for organic aquaculture applications. Technological advancements in aquaculture further complement these approaches. For example, Wang et al. (2021) developed a dual-stream 3D convolutional neural network (DSC3D) that integrates RGB and optical flow data from video clips to automatically recognise fish school behaviours, achieving high classification accuracy. Similarly, Yavuzer and Köse (2022) applied convolutional neural networks (CNNs) to predict fish quality through image analysis, highlighting the expanding role of artificial intelligence (AI) in monitoring fish health and product quality. Our study contributes to this growing body of knowledge by successfully applying behavioural data and machine learning for the non-invasive prediction of spawning events, demonstrating a novel application of AI to reproductive management in aquaculture, building on our previous work (Qadir et al., 2025). Machine learning approaches have also been applied to classify reproductive stages in fish based on morphological and physiological parameters (Flores et al., 2024; Yari et al., 2024), but these methods typically require handling and sampling of individuals.

In summary, this study addresses the multifaceted challenges of reproductive management in captive Senegalese sole and presents hormone-free IVF coupled with behavioural monitoring as a viable, less invasive alternative to hormone-based methods. Optimising sperm parameters and male selection remains critical to improving fertilisation outcomes, whilst behavioural analysis provides a powerful tool for predicting spawning readiness. Despite the promising outcomes of our hormone-free IVF approach and predictive model, some limitations remain. Variability in female gamete quality, timing sensitivity, and environmental factors continue to affect fertilisation success. Additionally, whilst our model demonstrated strong predictive performance, further validation with larger datasets and diverse conditions is needed to ensure robustness and broader applicability. Future research should focus on refining hormone-free fertilisation protocols, enhancing behavioural detection algorithms, and integrating these methods to advance sustainable and welfare-friendly aquaculture practices.

## 5. Conclusion

The findings of this study demonstrate that hormone-free fertilization is technically viable for Senegalese sole, with successful fertilization rates reaching 44% and hatching rates of 18% achieved using males selected for superior sperm quality. These results confirm that organic aquaculture protocols can produce viable offspring without hormonal intervention. However, the practical challenge lies not in the fertilization technique itself but in the timing: ovulated eggs remain viable for only approximately 3 hours, requiring precise identification of the ovulation event. To address this critical timing constraint, our automated behavioural monitoring system provides the necessary predictive capability. By detecting RTH and LA behaviours and achieving 82-85% accuracy in predicting ovulation nights, the system transforms hormone-free fertilization from an opportunistic process into a reliable, manageable protocol suitable for commercial implementation. This integration of targeted male selection, egg quality optimization, and behaviour-based ovulation prediction offers a comprehensive, non-invasive solution for reproductive management in flatfish aquaculture. The combined approach reduces dependence on hormonal treatments while maintaining the operational reliability required for commercial aquaculture, thereby supporting organic certification and promoting sustainable, ethical production systems. The automated prediction system is not merely an enhancement but an essential component that makes hormone-free protocols practically viable, enabling aquaculture managers to prepare resources and time interventions appropriately. This work demonstrates that machine learning applications, when properly integrated with biological understanding and practical constraints, can provide solutions that advance both sustainability goals and commercial viability in modern aquaculture.

## Acknowledgements

The study was funded by the European Union’s Programme H2020, project NewTechAqua GA862658, the Spanish project INIA – FEDER (RTA2014-00048) and the project PID2022-138327OB-I00 financed by the Ministerio de Ciencia e Innovación (MCIN)/Agencia Estatal de Investigación (AEI)/10.13039/501100011033/FEDER, UE. The participation of Abdul Qadir was supported by a Marti-Franquès 2020MFP-COFUND-18 EU Fellowship.

## Reference

Anguis, V., Cañavate, J.P., 2005. Spawning of captive Senegal sole (Solea senegalensis) under a naturally fluctuating temperature regime. Aquaculture 243, 133–145. 10.1016/j.aquaculture.2004.09.026

APROMAR, 2022. A guide on fish welfare in Spanish aquaculture - Volume 1: Concepts and generalities. Spanish Aquaculture Business Association.

APROMAR, 2019. Aquaculture in Spain. Spanish Aquaculture Business Association.

Beirão, J., Soares, F., Herráez, M.P., Dinis, M.T., Cabrita, E., 2009. Sperm quality evaluation in Solea senegalensis during the reproductive season at cellular level. Theriogenology 72, 1251–1261. 10.1016/j.theriogenology.2009.07.021

Cabrita, E., Soares, F., Dinis, M.T., 2006. Characterization of Senegalese sole, Solea senegalensis, male broodstock in terms of sperm production and quality. Aquaculture 261, 967–975. 10.1016/j.aquaculture.2006.08.020

Carazo, I., Chereguini, O., Martín, I., Huntingford, F., Duncan, N., 2016. Reproductive ethogram and mate selection in captive wild Senegalese sole (Solea senegalensis). Spanish Journal of Agricultural Research 14, e0401.

Du, L., Lu, Z., Li, D., 2022. Broodstock breeding behaviour recognition based on Resnet50-LSTM with CBAM attention mechanism. Computers and Electronics in Agriculture 202, 107404.

Duncan, N., Carazo, I., Chereguini, O., Mañanós, E., 2019. Mating Behaviour, in: Muñoz-Cueto, J.A., Sánchez, E.L.M., Vázquez, F.J.S. (Eds.), The Biology of Sole. CRC Press, Boca Raton, FL: CRC Press, Taylor & Francis Group, [2019] | “A science publishers book.,” pp. 169–184.

Fatsini, E., González, W., Ibarra-Zatarain, Z., Napuchi, J., Duncan, N.J., 2020. The presence of wild Senegalese sole breeders improves courtship and reproductive success in cultured conspecifics. Aquaculture 519, 734922.

García-López, A., Anguis, V., Couto, E., Canario, A.V.M., Cañavate, J.P., Sarasquete, C., Martínez-Rodríguez, G., 2006. Non-invasive assessment of reproductive status and cycle of sex steroid levels in a captive wild broodstock of Senegalese sole Solea senegalensis (Kaup). Aquaculture 254, 583–593. 10.1016/j.aquaculture.2005.10.007

González-López, W.Á., Ramos-Júdez, S., Giménez, I., Duncan, N.J., 2020. Sperm contamination by urine in Senegalese sole (Solea senegalensis) and the use of extender solutions for short-term chilled storage. Aquaculture 516, 734649.

Kotsiantis, S.B., Zaharakis, I.D., Pintelas, P.E., 2006. Machine learning: a review of classification and combining techniques. Artificial Intelligence Review 26, 159–190.

Martín, I., Carazo, I., Rasines, I., Rodríguez, C., Fernández, R., Martínez, P., Norambuena, F., Chereguini, O., Duncan, N., 2020. Reproductive performance of captive Senegalese sole, Solea senegalensis, according to the origin (wild or cultured) and gender. Spanish Journal of Agricultural Research 17, e0608.

Martín, I., Rasines, I., Gómez, M., Rodríguez, C., Martínez, P., Chereguini, O., 2014. Evolution of egg production and parental contribution in Senegalese sole, Solea senegalensis, during four consecutive spawning seasons. Aquaculture 424–425, 45–52.

Morais, S., Aragão, C., Cabrita, E., Conceição, L.E.C., Constenla, M., Costas, B., Dias, J., Duncan, N., Engrola, S., Estevez, A., Gisbert, E., Mañanós, E., Valente, L.M.P., Yúfera, M., Dinis, M.T., 2016. New developments and biological insights into the farming of *Solea senegalensis* reinforcing its aquaculture potential. Reviews in Aquaculture 8, 227–263. 10.1111/raq.12091

Porta, J., María Porta, J., Martínez-Rodríguez, G., Del Carmen Alvarez, M., 2006. Development of a microsatellite multiplex PCR for Senegalese sole (Solea senegalensis) and its application to broodstock management. Aquaculture 256, 159–166.

Qadir, A., Duncan, N., González-López, W.Á., Fatsini, E., Serratosa, F., 2025. Automated prediction of spawning nights using machine learning analysis of flatfish behaviour. Smart Agricultural Technology 12, 101668. 10.1016/j.atech.2025.101668

Ramos-Júdez, S., González-López, W.Á., Huayanay Ostos, J., Cota Mamani, N., Marrero Alemán, C., Beirão, J., Duncan, N., 2021. Low sperm to egg ratio required for successful *in vitro* fertilization in a pair-spawning teleost, Senegalese sole (*Solea senegalensis*). Royal Society Open Science 8, rsos.201718, 201718.

Rasines, I., Gómez, M., Martín, I., Rodríguez, C., Mañanós, E., Chereguini, O., 2012. Artificial fertilization of Senegalese sole (Solea senegalensis): Hormone therapy administration methods, timing of ovulation and viability of eggs retained in the ovarian cavity. Aquaculture 326–329, 129–135.

Rurangwa, E., Kime, D.E., Ollevier, F., Nash, J.P., 2004. The measurement of sperm motility and factors affecting sperm quality in cultured fish. Aquaculture 234, 1–28. 10.1016/j.aquaculture.2003.12.006

Salas-Leiton, E., Anguis, V., Martín-Antonio, B., Crespo, D., Planas, J.V., Infante, C., Cañavate, J.P., Manchado, M., 2010. Effects of stocking density and feed ration on growth and gene expression in the Senegalese sole (Solea senegalensis): Potential effects on the immune response. Fish & Shellfish Immunology 28, 296–302. 10.1016/j.fsi.2009.11.006

Shinde, P.P., Shah, S., 2018. A Review of Machine Learning and Deep Learning Applications, in: 2018 Fourth International Conference on Computing Communication Control and Automation (ICCUBEA). IEEE, Pune, India, pp. 1–6.

