## Supplementary Table S1 for "*In vitro* fertilisation procedure assisted with computer vision models for organic Senegalese sole (*Solea senegalensis*) culture"

Table 1. Sperm quality parameters assessed across 51 males classified into three groups: Low quality sperm (n = 24), High quality sperm (Not used) (n = 12), and High quality sperm used for IVF (n = 15). One-way ANOVA revealed significant differences among groups (F(2,48) = 23.36, P < 0.001). Tukey's post-hoc test showed that the High quality sperm used for IVF group had significantly higher motility than both the High quality sperm (Not used) group (P < 0.001) and the Low quality sperm group (P < 0.001), while no significant difference was found between the High quality sperm (Not used) and Low quality sperm groups (P = 0.469).

| **Parameter** | **Low quality sperm** | **High quality sperm (Not used)** | **High quality sperm used for IVF** | **High quality sperm used for IVF (After 24 hours)** |
| --- | --- | --- | --- | --- |
| **Motility** | 36.8 ± 14.5^a^ | 42.9 ± 16.4^a^ | 69 ± 13.2^b^ | 47 ± 17.4^a^ |
| **Concentration** | 12.0 ± 15.3 | 18.0 ± 19.9 | 28.3 ± 26 | 31 ± 24.30 |
| **VCL** | 47.3 ± 24.7 | 39.5 ± 20.6 | 80 ± 28.6 | 41.7 ± 18.7 |
| **VAP** | 32.3 ± 21.9 | 25.5 ± 19.2 | 64 ± 30 | 28 ± 17 |
| **VSL** | 24.3 ± 20.8 | 18.9 ± 17.7 | 55.4 ± 29.2 | 21 ± 15.6 |
| **STR %** | 55.2 ± 10.9 | 55.1 ± 9.3 | 69.1 ± 9 | 56.6 ± 9.6 |
| **LIN %** | 33.5 ± 13.4 | 31.1 ± 12.7 | 51.0 ± 13.0 | 33.5 ± 12.3 |
| **WOB %** | 56.5 ± 10.5 | 51.9 ± 11.7 | 67.3 ± 9.6 | 55.2 ± 10.4 |
| **ALH** | 1.1 ± 0.4 | 1.0 ± 0.4 | 1.3 ± 0.2 | 1.0 ± 0.2 |
| **BCF** | 5.6 ± 3.5 | 4.6 ± 2.2 | 11.1 ± 3.4 | 6.0 ± 4.0 |
